# Supramodal Coding of Detection and Confidence in the Human Cortex

**DOI:** 10.64898/2026.10.16.759637

**Authors:** Perrine Porte, Alexis Robin, Flore Boscher, Childéric Dezier, Dorian Goueytes, Shiva Mahdian, Dominique Hoffmann, Philippe Kahane, Lorella Minotti, Michael Pereira, Jeanne Caron-Guyon, Louise Goupil, Nathan Faivre

## Abstract

Perception is inherently multisensory, yet it remains unclear whether the brain detects stimuli and evaluates confidence in these detections using circuits shared across senses or circuits specific to each modality. Prior evidence is limited: scalp electrophysiology cannot localize where a shared code operates, whereas functional imaging cannot resolve its timing. We recorded intracranial activity (stereoelectroencephalography) in eight patients with epilepsy who performed a near-threshold audiovisual detection task in which they reported, on each trial, whether a visual, auditory, or audiovisual stimulus was present and rated their confidence, both overall and separately for each modality. Analyzing high-gamma activity at the single-channel and region-of-interest levels, we found that stimulus detection involved a broadly distributed, largely supramodal set of cortical regions responding regardless of modality, including the insula, dorsomedial prefrontal cortex, and superior temporal gyrus. Multivariate decoders trained to distinguish detected from undetected stimuli in one modality generalized to the other modality, indicating a shared detection code within these regions. Confidence relied on a more mixed architecture but was also substantially supramodal. Finally, the auditory cortex encoded confidence information about its non-preferred (visual) modality, with a similar effect in the visual cortex for auditory confidence, suggesting that the involvement of sensory cortices in confidence is not confined to its preferred input.

## 1. Introduction

Decades of work have mapped how the brain evaluates near-threshold sensory signals within modality-specific cortex, and identified regions where visual and auditory evidence converge (e.g., Noppeney et al., 2010). Yet, whether the brain relies on modality-specific regions or on shared, supramodal circuits to detect a stimulus and evaluate confidence in that detection remains largely unknown.

Two broad architectures can be considered. On one account, detection and confidence are assumed to rely primarily on modality-specific circuits. This view is grounded in the fact that signatures of stimulus detection are classically modality-specific: visual detection is reliably marked by the visual awareness negativity (VAN), an early negative-going event-related potential (ERP) component over occipital sites (Koivisto & Revonsuo, 2010). An analogous auditory awareness negativity (AAN) has been described over temporal sites (Eklund & Wiens, 2019). By directly comparing the two modalities in the same participants, Filimonov et al. (2022) confirmed this dissociation. At the metacognitive level, however, direct evidence for modality-specific confidence remains scarce. Most evidence for domain-specific metacognition instead comes from dissociations between perception and other cognitive domains, such as memory (Baird et al., 2013; Morales et al., 2018), rather than between sensory modalities within perception itself.

On an alternative account, detection and confidence are assumed to be governed by a shared, supramodal system regardless of sensory origin. At the perceptual level, this account is supported by multivariate decoding: Sanchez et al. (2020) showed that multivariate MEG correlates of stimulus detection in one modality predicted detection in another, and Niedernhuber et al. (2022) similarly reported overlapping markers of stimulus detection across visual, auditory, and somatosensory stimuli. At the metacognitive level, the supramodal view is supported by behavioral findings suggesting that confidence estimation is itself supramodal, with correlation in metacognitive performance across sensory modalities within individuals (Ais et al., 2016; de Gardelle et al., 2016), and confidence-related ERP amplitude showing a main effect of confidence with no interaction with modality (Faivre et al., 2018). A supramodal confidence signal has also been identified with fMRI during probabilistic inference, where confidence-related activity in parietal and prefrontal regions generalized across auditory and visual sessions (Meyniel & Dehaene, 2017), and recent scalp-EEG work has shown a decision-related signal that generalizes across modalities to predict confidence (Davidson et al., 2025), paralleling supramodal decision signals in perceptual decision-making (O’Connell et al., 2012). This domain-general account of confidence, however, has been established almost entirely using discrimination or inference tasks (e.g., Ais et al., 2016; de Gardelle et al., 2016; Faivre et al., 2018; Meyniel & Dehaene, 2017). Whether domain-generality extends to detection, a computationally distinct judgment (Mazor et al., 2020; Mazor et al., 2025; Porte et al., 2026), remains an open question. Anatomically, the two accounts diverge: modality-specific processing predicts effects confined to sensory cortices (occipital and superior temporal gyrus), whereas a supramodal account predicts involvement of amodal association and monitoring regions, including the insula and frontal regions.

To our knowledge, no study has directly decoded a shared neural marker for confidence across modalities during detection. In the current preregistered study, we used stereotactic electroencephalography (sEEG), which provides the temporal resolution required for cross-modal decoding, as well as direct access to deep brain structures and high-gamma activity – which has been shown to reflect local neuronal firing (Nir et al., 2007; Lachaux et al., 2012). Patients with epilepsy implanted with intracranial electrodes for clinical purposes performed a near-threshold detection task in which they judged whether a visual, auditory, or audiovisual stimulus was present, regardless of modality, and rated their confidence. Our paradigm also leverages a bidimensional confidence scale, on which participants reported their detection and their confidence for each modality simultaneously. Combining this paradigm with sEEG, we tested, at the level of individual channels and anatomically defined regions of interest, whether high-gamma activity reflects detection and confidence in a modality-specific or supramodal fashion, notably by testing whether decoders trained on one modality generalize to the other.

## 2. Methods

### 2.1. Participants and sEEG data collection

Twelve participants (three self-reported women, mean age = 31.58 (*SD* = 10.11), see Table S1) with drug-resistant focal epilepsy were recruited at the Grenoble Alpes University Hospital. They were on the following antiseizure medications: carbamazepine (5), perampanel (2), clobazam (2), levetiracetam (1), lamotrigine (1), cenobamate (3), lacosamide (1). In addition, one participant was on antidepressants (sertraline). All patients gave informed consent before taking part in the experiment, which was approved by the ethics committee Sud-Ouest et Outre-Mer 4 (2017-A03248-45).

They were implanted with semi-rigid linear stereotactic electrodes (DIXI Medical, France, Alcis France), and data were recorded with a sampling rate of 512Hz using an Audio-Visio Micromed recording system (Treviso, Italy). The anatomical locations of electrode contacts, decided uniquely by clinical purpose, were identified on BrainVisa using a post-implantation CT scan co-registered with a pre-implantation T1 MRI scan.

Four participants were excluded from all analyses because their detection accuracy was nearly zero in either the auditory and/or the visual modality. Three of the remaining eight participants did not complete the entire experiment due to clinical constraints (out of six blocks, two completed four blocks and one completed three blocks).

### 2.2. Protocol

The experimental paradigm consisted of an audiovisual detection task, similar to that of Porte et al. (2026). It was programmed and presented using PsychoPy v2023.2.3. The visual stimulus was a light grey circle with a diameter of 14mm embedded in dynamic visual Gaussian noise updated every 33 ms (30 Hz). The auditory stimulus was a 1kHz sinusoidal tone embedded in auditory pink noise. The sound was played through air-conduction earphones (ER3C Insert Earphones, Etymotic, USA). On each trial, the stimulus could be visual, auditory, audiovisual, or absent. Participants were asked to judge whether a stimulus was present regardless of its modality. After this amodal detection response, participants indicated their confidence in their answer on a continuous scale from 0% (sure incorrect) to 100% (sure correct). Finally, on a bidimensional (audio/visual) scale, they indicated simultaneously their detection response (present/absent) and confidence for both modalities (Fig.1).

**Figure 1:**
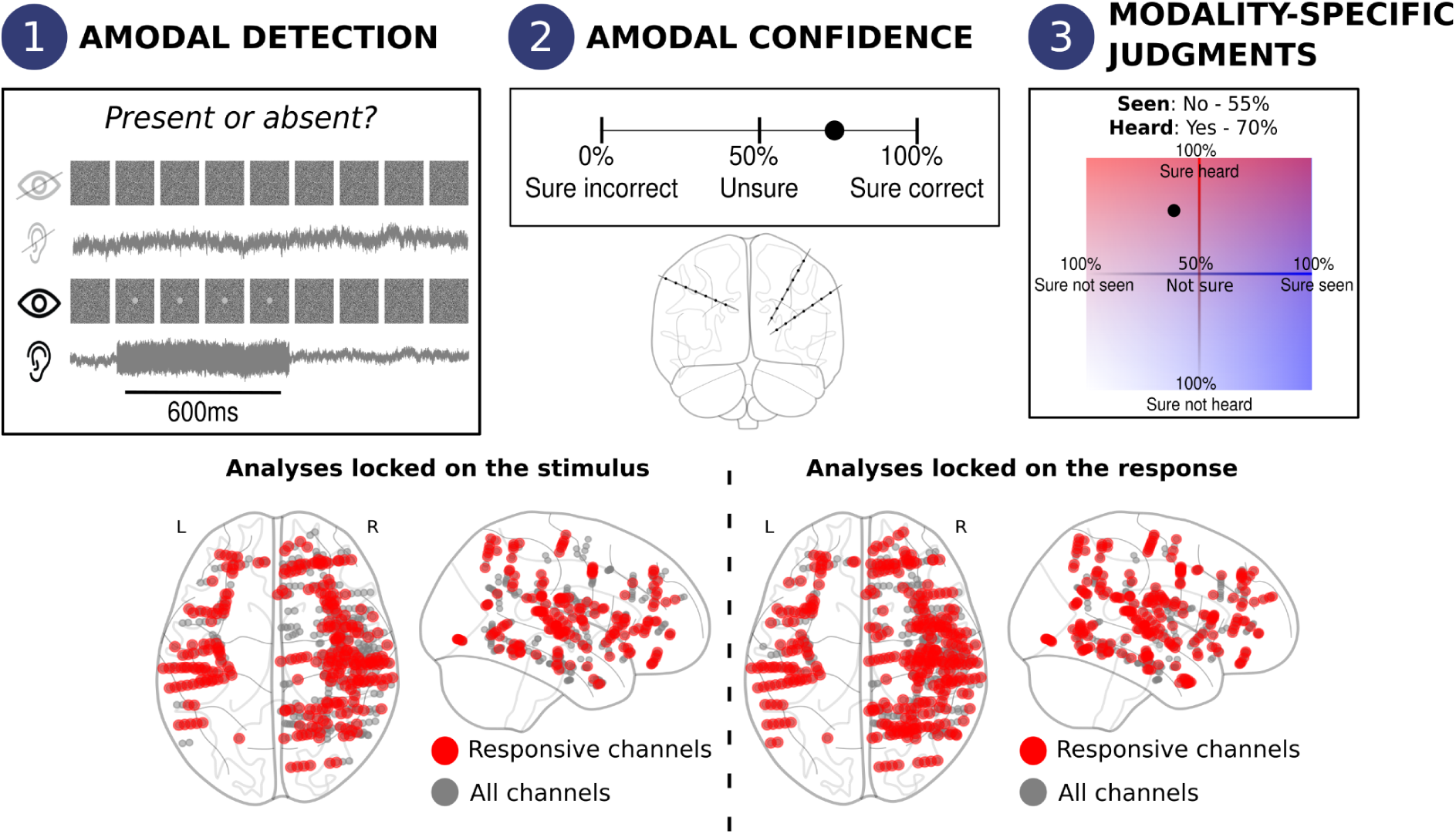
Top: Experimental paradigm. 1) Amodal detection irrespective of the stimulus modality. Representation of the different possible conditions: visual stimulus absent, auditory stimulus absent, visual stimulus present, auditory stimulus present; 2) Amodal confidence in the detection response; 3) Modality-specific judgments: Simultaneous detection and confidence answer for each modality (i.e., auditory and visual). At the same time, we recorded intracranial activity from patients with epilepsy implanted with stereotactic electrodes. **Bottom: Channel localisation.** All analyzed channels are depicted in grey. All responsive channels are depicted in red, for both stimulus-locked and response-locked analyses.

Stimulus intensity was calibrated for each participant at the beginning of the experiment to achieve a 50% hit rate in the visual and auditory conditions based on unimodal psychometric curves. Auditory and visual calibrations were conducted separately, and their order of presentation was counterbalanced across participants. To estimate each psychometric curve, we asked participants to detect stimuli at five different intensities, each presented 20 times, with 20 additional trials where no stimulus was presented. This calibration phase was repeated if the psychometric curve fitting did not converge. As revealed by a pilot study, stimulus detectability increased between the calibration and the main experimental phase. To account for this increase, the stimulus intensity for the main experiment was preregistered as the intensity corresponding to 40% of detected stimuli during the calibration phase.

Once calibration was complete, participants performed the main task. Each trial began with the presentation of audiovisual noise. The stimulus could appear 200, 300, or 400ms after the noise onset and lasted 600ms. In the audiovisual condition, the visual and auditory stimuli were always synchronous. Participants indicated whether a stimulus was present or absent, irrespective of the perceived modality, by pressing the right or left arrow on the keyboard. The response keys associated with each response were counterbalanced across participants. Participants could respond (*amodal detection answer*) immediately from the start of the trial, with a 4s time limit after the stimulus offset. If they did not respond during this time period, a timeout message appeared, and they had to restart a trial. With a 100ms delay after the response, the amodal confidence scale was presented until a response was provided using the mouse. Immediately after, the bidimensional (audio/visual) scale was presented with each axis ranging from 100% (sure not perceived) to 100% (sure perceived).

After 10 training trials, participants completed 6 experimental blocks of 48 trials, resulting in a total of 288 trials. Within each block, experimental conditions were equiprobable and presented at random.

### 2.3. Behavioral analysis

All behavioral statistical analyses were performed in R (version 4.5.2, main packages used: *tidyverse* (v2.0.0, Wickham et al., 2019), *lme4* (v1.1.38, Bates et al., 2015), *ggplot2* (v4.0.1, Wickham, 2016), *psycho* (v0.6.1, Makowski, 2018)), following our preregistered plan (https://osf.io/cfdvq/overview).

We calculated individual d-primes and response criteria to quantify, respectively, stimulus detectability and bias toward reporting that the stimulus was present or absent. To investigate amodal detection performance and metacognitive sensitivity, we conducted a mixed-effects logistic regression on accuracy as a function of the stimulus modality (auditory, visual, audiovisual, or absent), amodal confidence, and their interaction. We also fitted a mixed-effects logistic regression on the accuracy of absence judgments (misses vs. correct rejections) as a function of amodal confidence, to assess participants’ ability to monitor their absence judgments. To investigate confidence bias, we fitted two linear models: one comparing the mean amodal confidence as a function of stimulus modality, and the other comparing the mean amodal confidence as a function of participants’ judgments (stimulus judged absent or present). Finally, we estimated metacognitive efficiency using response-conditional hierarchical Mratio (Hmeta-d’, Fleming, 2017), binning the confidence into six bins based on the overall confidence distribution of each participant.

At the modality-specific level, we computed four models for each modality. Each model examined how confidence in a given modality interacted with the presence or absence of a stimulus in the other modality, separately for trials in which the stimulus was physically present or absent in the given modality, and for trials in which it was judged as present or absent.

- *accuracy in detecting presence (hit vs miss) ∼ modality-specific confidence * presence in the other modality;*
- *accuracy in detecting absence (false alarms [FA] vs correct rejection [CR]) ∼ modality-specific confidence * presence in the other modality;*
- *absence judgment accuracy (miss vs CR) ∼ modality-specific confidence * presence in the other modality;*
- *presence judgment accuracy (hit vs FA) ∼ modality-specific confidence * presence in the other modality*.

Finally, we investigated the relationship between modality-specific and amodal confidence judgments by first fitting a linear model of mean confidence as a function of confidence type (i.e., amodal, auditory, or visual) to examine differences in confidence biases. Then, to examine the contribution of each modality to amodal confidence, we fitted a linear model of amodal confidence as a function of auditory and visual confidence.

### 2.4. sEEG preprocessing

All preprocessing steps were performed in Python (version 3.11.9) using *mne* (v1.6.1, Gramfort et al., 2013).

We first excluded channels localized outside the grey matter and those displaying epileptiform or artifactual activity, as determined by visual inspection. We then performed a bipolar re-referencing between two neighbors among the remaining channels; bipolar re-referencing between two channels was performed when a maximum of one channel was removed in between (e.g., if u2 was excluded, the bipolar re-referencing u1-u3 was authorized; but if u2 and u3 were excluded, the bipolar re-referencing u1-u4 was not performed). Only the bipoles in which the two individual channels were located in the same anatomical region based on the MarsAtlas parcellations (Auzias et al., 2016) were kept.

We then extracted the high-gamma activity (HGA) between 70 and 150 Hz for these bipoles. We divided this frequency range into seven half-overlapping frequency bands of 20Hz each (i.e., 70-90Hz, 80-100Hz, etc.). For each band, the signal was split into epochs, from −1 to 2s around the stimulus onset and from −6 to 1s around the response. For each band, each trial was baseline-corrected (i.e., each trial was normalized by the mean baseline across trials), with a baseline from −0.2s to stimulus onset for analyses locked to stimulus onset, and from 0 to 0.2s into the trial for analyses locked to the response. Then the different bands were averaged. Individual epochs were then visually inspected to remove remaining artefactual or epileptiform activity. Finally, the data were downsampled to 64Hz to minimize the number of statistical tests while preserving meaningful temporal resolution.

### 2.5. Trial Segmentation and Channel Selection

Statistical analyses were performed on a time window from −0.1 to 0.7s around the stimulus onset and from −0.5 to 0.5s around the response.

Channel selection for each participant was performed separately for stimulus-locked and response-locked analyses. For stimulus-locked analyses, a channel was considered responsive if average high-gamma activity showed a significant difference between a pre-stimulus window (−0.2 to 0s relative to stimulus onset) and a post-stimulus window (0 to 1s after stimulus onset; paired Wilcoxon test). Of 580 channels, 274 (i.e., 47%) were found to be responsive. Similarly, for response-locked analyses, a channel was considered responsive if it showed a significant difference between a baseline window (first 0.2s of each trial, before stimulus onset) and a peri-response window (−0.5 to 0.5s around the response onset; paired Wilcoxon test). Of 580 channels, 333 (i.e., 57%) were found to be responsive (Fig.1).

### 2.6. Statistical Analysis of High-Gamma Activity

All statistical analyses were performed in Python, using *numpy* (v1.26.4), *pandas* (v2.2.2), *scipy* (v1.13.0), and *pymer4* (v0.8.0). We fitted generalized linear models on HGA among responsive channels. Models were applied to each time point within the analysis window, and a time-wise false-discovery rate (FDR) correction procedure was applied to correct for multiple comparisons. Only significant effects lasting at least 50ms were kept.

To assess deviation from chance level, we performed prevalence permutation tests with 1000 permutations. For each of our by-channel models, we tested whether the number of channels with a significant effect exceeded chance within each ROI. For each permutation, for every responsive channel within an ROI, we shuffled the labels of each of our models’ effects. The permuted p-value was computed as (permutations with count >= observed + 1) / (number of permutations + 1).

Some participants were excluded from specific analyses due to a lack of confidence variability, or a small number of trials in one experimental condition. The number of participants included in each analysis is available in the SI (Table S2).

All the models described below were first applied to each responsive channel. Individual responsive channels were then grouped by ROI across participants and submitted to the same models, including nested random intercepts by participant and channel. Analyses at the ROI level were performed on regions containing at least 20 channels.

#### 2.6.1. Analysis Locked on the Stimulus Onset

We first fitted a model on HGA as a function of each main effect of interest: the stimulus modality (visual, audio, audiovisual), the response accuracy (hit vs miss), and the amodal confidence. We defined detection-specific channels as those showing a main effect of response accuracy. Then, we computed the interaction between response accuracy and stimulus modality, to determine whether detection-specific channels were amodal (difference present both for visual and auditory trials), unimodal (difference present only for the auditory or visual trials), or multimodal (difference larger for audiovisual compared to unimodal trials). We also computed the interaction between stimulus modality and amodal confidence for all trials, and then separately for correct and incorrect responses.

These models were extended to modality-specific responses at the visual and auditory levels. One model investigated the interaction between stimulus modality and response accuracy, and another investigated the interaction between stimulus modality and confidence, for all trials and then separately for correct and incorrect responses.

Finally, we examined the link between amodal and modality-specific responses. We investigated whether there was a general effect of confidence, irrespective of its type (i.e., auditory, visual, or amodal), and whether it differed as a function of confidence type. The same was done with response accuracy. Note that the model investigating the link between the different accuracy responses was not preregistered.

ROI-level analyses were conducted on the left inferior parietal cortex (23 responsive channels), the right insula (21 responsive channels), the right superior temporal gyrus (35 responsive channels), and the right visual cortex (28 responsive channels).

#### 2.6.2. Analysis Locked on the Detection Response

The same models as in the stimulus-locked analyses were fitted to response-locked HGA. We additionally fitted a model that included the judgment (i.e., whether the stimulus was judged present or absent) and its interaction with amodal confidence, separately for all trials and for correct responses only. Finally, focusing on absence judgments, we fitted a model that included the interaction between amodal confidence and response accuracy (CR vs. miss).

ROI-level analyses were conducted in the left inferior parietal cortex (20 responsive channels), the right superior temporal gyrus (36 responsive channels), and the right visual cortex (48 responsive channels).

### 2.7. Multivariate Analysis

We performed multivariate analyses locked on the stimulus onset using the *sklearn* package (v1.4.2). Namely, we tested the transferability of LDA decoders trained on detection across modalities (e.g., training on hits vs. misses for auditory stimuli and testing on hits vs. misses for visual stimuli). Similarly, we tested the transferability of LDA decoders trained on binarized modality-specific confidence (e.g., training on auditory confidence and testing on visual confidence), with confidence binned into two equal-width bins. To avoid the accuracy confounding factor, the decoder applied to confidence was only applied to correct answers.

These decoders were applied separately to each participant on ROIs of at least five channels. A total of 13 ROIs were found across all participants (see SI.1 for the complete list).

We applied each decoder with a sliding window of 150ms and a step size of 20ms, resulting in a time-course of decoding accuracy across windows. We performed cross-validation with 5 folds, using stratified folds to have the same proportion of each class in each fold. Each fold was split into training and test sets. To address unbalanced design in the training data, we performed random undersampling of the more frequent category and SMOTE-based data augmentation of the less frequent category. Hyperparameter optimization (i.e., solver and shrinkage parameters) was performed using nested cross-validation, with AUROC as the optimization metric. These optimization steps were done on the training modality. To test the decoders’ ability to generalize across modalities, the final optimized model was trained on the entire dataset of one modality and tested on the other. We used permutation tests to estimate decoding performance: we trained the same decoder with the same hyperparameters on data with shuffled labels, performing 1000 permutations and estimating p-values from these permutations.

### 2.8. Deviations from preregistration

Our study followed the preregistered plan (https://osf.io/cfdvq/overview) except for the following deviations, which we report here for transparency. We planned to stop collecting data after a minimum of 15 exploitable datasets. However, within the data collection period – the data collection started in spring 2024, and had to end at the end of spring 2026 because of the timeline of the first author’s PhD – we were unable to reach this number. At the behavioral level, the mapping of the auditory and visual modalities to the horizontal and vertical axes of the bidimensional confidence scale was counterbalanced across participants to avoid response and motor biases. Regarding neural analyses, we preregistered a minimum of 25 task-responsive channels per region for ROI-level analysis. Because of the limited sample size, we lowered this threshold to 20 channels and regrouped the original MarsAtlas parcels into larger anatomical ROIs, which were validated by a trained neurologist (see SI.2 for the complete regrouping). For stimulus-locked analyses, we defined the pre-stimulus baseline window as −0.2 to 0s relative to stimulus onset, rather than the preregistered −0.5 to −0.1s window, which would have included the audiovisual noise onset, thereby contaminating the baseline with evoked activity. Also given our limited sample size, we selected responsive channels using uncorrected Wilcoxon tests rather than permutation tests to retain as many channels as possible. Nevertheless, we additionally computed unpreregistered prevalence permutation tests for every by-channel model.

### 2.9. Code Accessibility

Behavioral data, experimental protocol, and analysis code are available at https://gitlab.com/nfaivre/bimodal_confidence_seeg_public. High-gamma activity data are available on the OpenNeuro database (https://doi.org/10.18112/openneuro.ds008959.v1.0.0)

## 3. Results

### 3.1. Behavioral Results

At the perceptual level, participants detected stimuli with above-chance sensitivity (*amodal d’* = 2.49 (*SD* = 0.61); *modality-specific auditory d’ =* 2.38 (*SD* = 0.44); *modality-specific visual d’* = 2.04 (*SD* = 0.83), Fig.2A). The logistic regression analyses on amodal accuracy showed that participants detected bimodal stimuli more accurately than unimodal ones (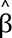 = 0. 42, 95% CI [0. 31, 0. 54], *z* = 7. 39, *p* <. 001) (Fig.2D). There was no difference in detection performance between auditory and visual stimuli, indicating that our performance matching across modality was efficient (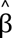 =− 0. 33, 95% CI [− 0. 81, 0. 15], *z* =− 1. 34, *p* =. 18). Additionally, participants had a significant bias toward responding that the stimulus was absent using the amodal scale (*M_criterion_* = 0. 61, 95% CI [0. 28, 0. 93], *t*(7) = 4. 38, *p* =. 003; Fig.2A). We confirmed this was also the case for each modality by deriving modality-specific detection responses using the modality-specific scale (**visual**: *M* = 1. 03, 95% CI [0. 54, 1. 52], *t*(7) = 4. 99, *p* = . 002; **auditory**: *M* = 0. 79, 95% CI [0. 44, 1. 13], *t*(7) = 5. 38, *p* = . 001) (see Table S3 for reaction times).

**Figure 2:**
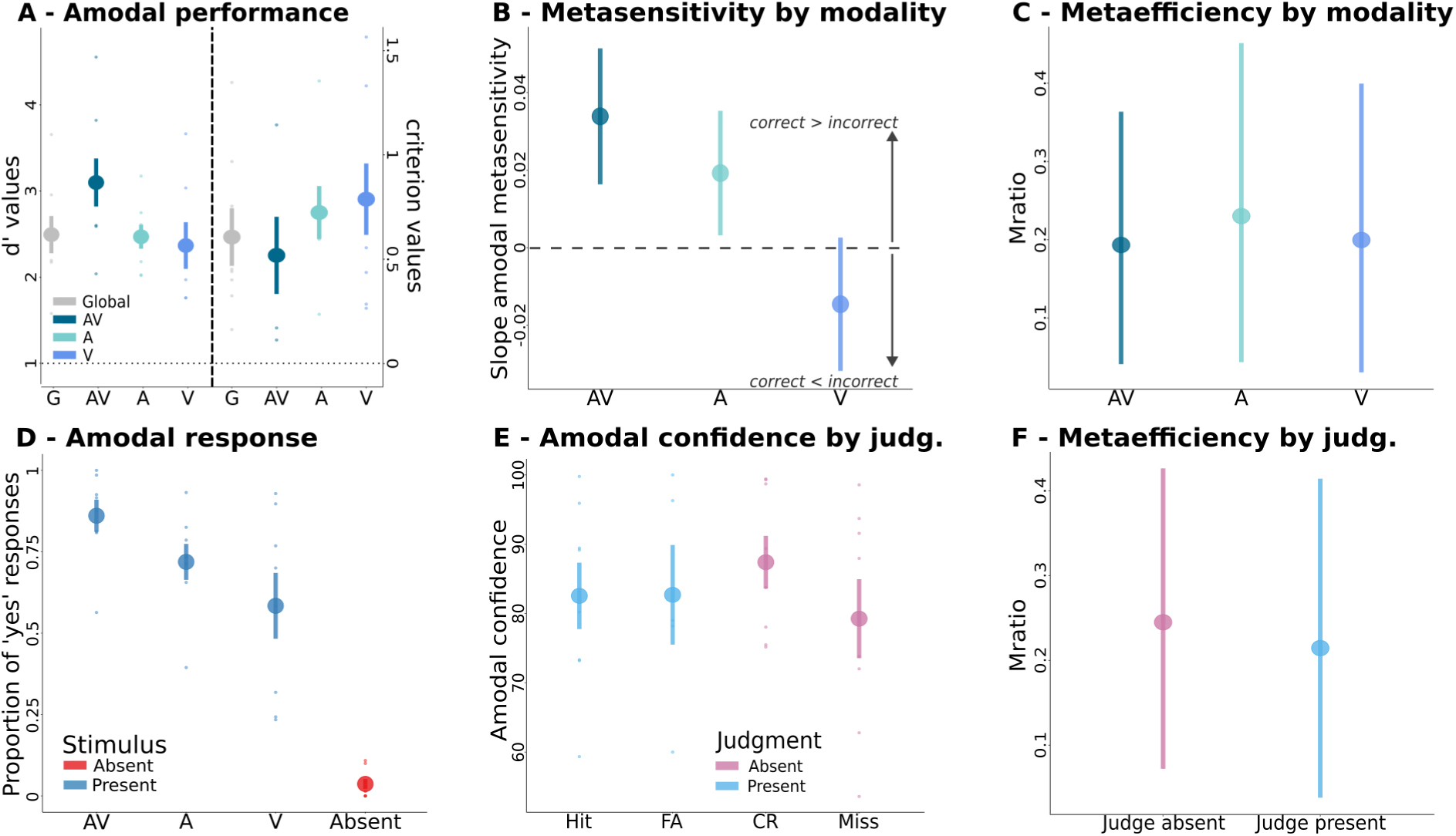
Behavioral results. A) Amodal d’ (left panel) and criterion (right panel), globally and by stimulus modality (Global: all experimental conditions together; AV: audiovisual; A: auditory; V: visual). B) Slopes of the logistic regression between accuracy and confidence as a function of the stimulus modality. C) Mratio estimation for presence judgments as a function of the stimulus modality. D) Proportion of stimuli judged present as a function of the experimental condition. E) Amodal confidence as a function of both accuracy and judgment type. F) Mratio estimation for presence and absence judgments.

At the metacognitive level, we did not observe a difference in amodal confidence between absence and presence judgments (Fig.2E), nor between stimulus modalities (*p* > .05). However, participants adapted their amodal confidence better to the accuracy of the response in bimodal than in unimodal trials (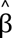 = 0. 01, 95% CI [0. 00, 0. 02], *z* = 2. 35, *p* =. 02) and in auditory than in visual trials (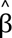 =− 0. 02, 95% CI [− 0. 03, 0. 00], *z* =− 2. 26, *p* =. 02) (Fig.2B). Moreover, participants reported higher amodal confidence in correct than in incorrect absence judgments (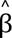 =− 0. 05, 95% CI [− 0. 09, − 0. 01], *z* =− 2. 71, *p* =. 007).

To evaluate metacognitive performance irrespective of task performance, we analyzed metacognitive efficiency (i.e., Mratio) for presence versus absence judgments, and across stimulus modalities. We found no credible difference in Mratio between presence and absence judgments (Δ*M* = −0.03 [-0.30, 0.25]; Fig.2F), nor as a function of stimulus modality (**bimodal vs unimodal trials**: Δ*M* = −0.02 [-0.32, 0.25]; **auditory vs visual trials**: Δ*M* = 0.03 [-0.27, 0.34]; Fig.2C).

At the modality-specific level, we did not observe any significant effect of the presence of a stimulus in the other modality on perceptual and metacognitive performance in a given modality (all *p* >.05, e.g., the presence of an auditory stimulus does not significantly increase the probability of judging that a visual stimulus was present).

Finally, we observed that participants tended to be more confident in their modality-specific judgments than in their amodal ones (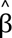 =− 1. 81, 95% CI [− 2. 86, − 0. 76], *t*(14) =− 3. 38, *p* =. 004); with no difference between auditory and visual confidence (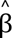 = 0. 06, 95% CI [− 1. 76, 1. 88], *t*(14) = 0. 07, *p* =. 95). Moreover, both the auditory and visual confidence predicted the amodal confidence (**auditory confidence**: 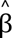 = 0. 45, 95% CI [0. 32, 0. 58], *t*(7. 08) = 6. 75, *p* <. 001; **visual confidence**: 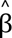 = 0. 37, 95% CI [0. 27, 0. 48], *t*(6. 28) = 6. 93, *p* <. 001)). Together, these behavioral results confirm that our paradigm successfully captured detection and metacognitive processes across modalities: participants detected bimodal stimuli more accurately than unimodal ones, and their confidence tracked response accuracy, including for absence judgments. On this basis, we turn to examine the neural markers of detection and confidence in presence and absence across modalities.

### 3.2. Role of Specialized Regions for Unisensory and Multisensory Perception

Across the results, we report single-channel effects only when the associated prevalence permutation test was significant (see SI.3 for regions not exceeding chance).

As expected, we found stimulus-evoked high-gamma responses in the right visual cortex (VC) and the right auditory cortex (i.e., the superior temporal gyrus, STG) in their preferred modality, at the single channel level (*N* = 7 channels in the right STG, ***permutation***: *p* <.001, *N* = 2 channels in the left STG, ***permutation***: *p* =.002, *N* = 6 channels in the right VC, ***permutation***: *p* <.001) and at the ROI level during a sustained period from 50ms to 700ms in the right STG and from 257ms to 700ms in the right VC (see SI for other regions).

Interestingly, these sensory-specific areas also showed stronger stimulus-evoked responses for bimodal than unimodal trials at the ROI level in both the right VC (significant effect of modality from 555ms to 700ms) and the right STG (significant effect of modality from 115ms to 398ms and from 445 to 586ms). These results corroborate the implication of specialized sensory cortices for multisensory perception.

### 3.3. Architecture of Detection Network

Having shown that high-gamma activity in sensory cortices increased following the presentation of unimodal and bimodal stimuli, we examined whether the activity reflecting detection accuracy depended on the stimulus modality.

#### 3.3.1. Activity Restricted to Modality-specific Detection

To determine activity specific to the modality of presentation, we determined which channels responded to the modality-specific detection accuracy (i.e., visual and auditory accuracy as defined by the bidimensional report scale). Furthermore, we tested whether channels showed a differentiated response to the amodal accuracy as a function of the stimulus modality, and whether channels responded specifically to one response type (i.e., amodal or modality-specific judgments).

Evoked high gamma responses differed between auditory correct and incorrect responses in the left somatosensory cortex (SSC; *N* = 1, *p* = .01) and right STG (*N* = 1, *p* = .04). For visual responses, one channel in the right orbitofrontal cortex (OFC; *N* = 1, *p* = .02) had differentiated activity as a function of visual accuracy. The effect of visual accuracy also differed between visual-only and audiovisual trials in the left and right dorsolateral prefrontal cortex (dlPFC; *N_left_* = 1, *p_left_* = .002; *N_right_* = 2, *p_right_* < .001). Finally, one channel in the left SSC (*p* = .02) was considered multimodal, with an interaction between response accuracy and modality (bimodal vs unimodal trials). One channel in the left inferior parietal cortex (IPC; *p* = .02) was modality-specific, showing an interaction between response accuracy and modality (visual vs auditory trials). We also note that no channels were found to respond differently to the modality of the detection response (i.e., amodal or modality-specific responses). Together, these results show limited cortical evidence for modality-specific encoding of stimulus detection.

#### 3.3.2. Supramodal Network of Detection

In contrast, we identified a large set of cortical regions reflecting amodal stimulus detection accuracy (i.e., regardless of sensory modality), including the left inferior frontal cortex (IFC; *N* = 3, *p* < .001), the left and right insula (*N_left_* = 2, *p_left_*= .005; *N_right_* = 2, *p_right_* = .01), the right motor cortex (MC; *N* = 2, *p* = .01), the right posterior cingulate cortex (PCC; *N* = 1, *p* = .04), and the right STG (*N* = 3, *p* = .01), with latencies spanning from 178ms to 610ms. We additionally identified regions in which accuracy effects were consistently observed whether accuracy was defined from amodal responses or separately for the auditory and visual responses on the modality-specific scale. This included the left and right IFC (*N_left_ =* 1*, p_left_* <.001; *N_right_ =* 1, *p_right_* = .02), left IPC (*N =* 1*, p* = .02), left and right insula (*N_left_ =* 2*, p_left_* < .001; *N_right_*= 1, *p_right_* = .03), left and right SSC (*N_left_ =* 1, *p_left_* = .02; *N_right_* = 1, *p_right_* = .01), right anterior cingulate cortex (ACC; N = 1, *p* = .008), right MC (*N =* 2, *p* = .002), right PCC (*N* = 1, *p* = .01), right superior parietal cortex (SPC; *N* = 1, *p* = .03), right STG (*N* = 4, *p* = .002), and right dlPFC (*N* = 1, *p* = .02), with latencies spanning from 50ms to 630ms (Fig.3A-B).

**Figure 3:**
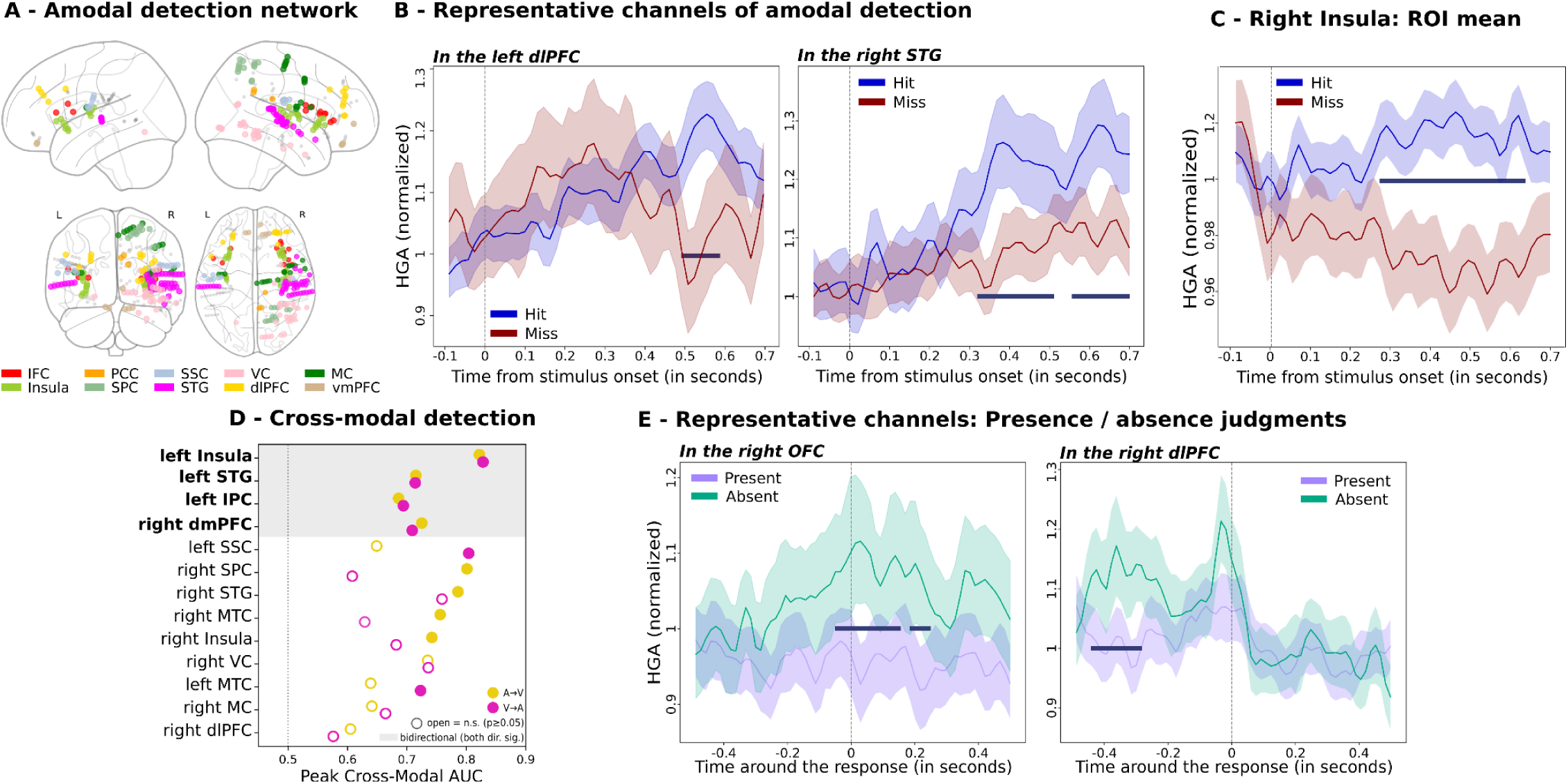
A distributed supramodal network supports stimulus detection. A) Cortical distribution of amodal detection effects regardless of stimulus modality. Channels whose high-gamma activity distinguishes detected (hit) from undetected (miss) stimuli irrespective of modality, plotted on a template brain and colored by region; all other responsive channels are shown in grey. B) High-gamma time courses for two representative significant channels (in the left dlPFC and right STG), showing separation of hits from misses; trial mean ± 95% CI. C) Region-level high-gamma in the right insula for hits and misses; mean ± 95% CI. D) Cross-modal detection decoding. Peak generalization AUC is shown for each region and transfer direction (orange: auditory→visual; pink: visual→auditory); filled symbols denote significant decoding (permutation *p* < .05), open symbols non-significant; the dashed line marks chance (AUC = 0.5), and the shaded band groups the four bidirectionally-decoding regions. E) High-gamma time courses for two representative significant channels (in the right OFC and right dlPFC), showing separation for presence and absence judgments; trials mean ± 95% CI.

Beyond single-channel analyses, analyses at the ROI level showed that the right insula seemed to play a central role in supramodal detection, as it responded consistently to accuracy, whether it was defined from amodal responses (amodal scale) or separately for the auditory and visual responses (bidimensional scale). The insula also responded to amodal accuracy regardless of stimulus modality (auditory, visual, or audiovisual) with a latency from 270ms to 630ms (Fig.3C). The only region in which a similar effect was found was the right STG, which was somewhat unexpected given that it has traditionally been associated with auditory processing (detection effect regardless of modality from 320 to 660ms).

While these univariate analyses identify regions where activity co-varies with detection, they cannot establish whether the same neural code underlies detection across modalities. To directly test this, we trained a decoder on the difference between hits and misses in one modality (e.g., for visual stimuli) and then assessed whether it could decode hits and misses in the other (e.g., for auditory stimuli). Decoding transfer from visual to auditory detection accuracy exceeded chance-level in the right dorsomedial prefrontal cortex (dmPFC; *AUC peak*: 0.71 at 60ms), left insula (*AUC peak*: 0.83 at 530ms), left IPC (*AUC peak*: 0.70 at 530ms), left SSC (*AUC peak*: 0.80 at 450 ms), left middle temporal gyrus (MTG; *AUC peak*: 0.72 at 250ms), and left STG (*AUC peak*: 0.71 at 330ms). Conversely, decoding transfer from auditory to visual detection occurred in the right dmPFC (*AUC peak*: 0.73 at 50ms), bilateral insula (**left**: *AUC peak*: 0.82 at 470ms; **right**: *AUC peak*: 0.74 at 530ms), left IPC (*AUC peak:* 0.69 at 30ms), bilateral STG (**right**: *AUC peak*: 0.79 at 150ms, **left**: *AUC peak*: 0.72 at 350ms), right MTG (*AUC peak*: 0.76 at 180ms), and right SPC (*AUC peak:* 0.80 at 50ms). Notably, decoding generalized bidirectionally in the right dmPFC, left insula, left IPC, and left STG, providing the strongest evidence for a symmetric detection code in these regions; generalization in the remaining regions was direction-specific, which may reflect true asymmetries in how each modality’s detection signal is represented, or differences in decoding sensitivity between the auditory and visual modalities (Fig.3D).

Altogether, these results point to a supramodal architecture for perceptual detection, with only sparse, scattered evidence of modality-specific coding.

#### 3.3.3. Are Presence and Absence Judgments Supported by Distinct Brain Regions

Having characterized neural correlates of objective detection accuracy, we asked whether cortical activity also differentiated trials as a function of participants’ subjective judgment (present versus absent). Amodal analyses were performed on high-gamma activity locked to the response, as they included trials with no stimulus.

At the single-channel level, high-gamma responses differentiating amodal presence and absence judgments were observed in the left and right IPC (*N_left_* = 2, *p_left_* = .02; *N_right_* = 2, *p_right_* = .01), the right MC (*N* = 3, *p* <.001), the left MTG (*N* = 2, *p* = .02), the left and right OFC (*N_left_* = 3, *p_left_* <.001; *N_right_*= 3, *p_right_* = .002), the left and right SSC (*N_left_* = 2, *p_left_* = .004; *N_right_* = 4, *p_right_* = .007), the right VC (*N* = 12, *p* < .001), the left ventromedial prefrontal cortex (vmPFC; *N* = 1, *p* = .03), the right IFC (*N* = 2, *p* = .006), the right SPC (*N* = 5, *p* < .001) and the right dlPFC (*N* = 2, *p* = .02). However, due to our lower number of incorrect presence judgments (on average across participants, they represent 4% of the trials), these results might be contaminated by the objective accuracy. (Fig.3E).

Focusing solely on correct responses, differences in HGA between absence (i.e., correct rejections) and presence judgments (i.e., hits) were found in the left and right OFC (*N_left_* = 1, *p_left_* = .03; *N_right_* = 2, *p_right_* = .002), the left VC (*N* = 1, *p* = .04), the right IFC (*N* = 2, *p* < .001), the right IPC (*N* = 2, *p* = .003), and the right dlPFC (*N* = 2, *p* = .003), with latencies spanning from −490ms to 440ms around the response (see SI for regions where prevalence did not exceed chance). Yet, these results could also reflect an effect of the objective stimulus presence.

To account for these limitations, we crossed the regions showing an effect in the two previous models to approximate the neural correlates of presence and absence judgments, revealing differentiated activity as a function of judgment in the left and right OFC, the right IFC, the right IPC, and the right dlPFC (Fig.3E).

### 3.4. A Mixed Modal and Supramodal Network for Confidence

We applied the same analysis strategy to confidence, examining whether the activity reflecting confidence ratings depended on the stimulus modality and on the type of confidence (i.e., amodal or modality-specific confidence). We report results from analyses performed both irrespective of detection accuracy and only on correct responses, both locked on the stimulus onset and on the response.

#### 3.4.1. Modal Areas for Confidence

Time-locked to the stimulus onset, activity differentiated auditory from visual confidence in the right MC (*N* = 2, *p* < .001) and the right STG (*N* = 1, *p* = .04). Modality-specific analyses showed that auditory confidence modulated activity in the right insula (*N* = 2, *p* < .001). For correct auditory responses, this effect was also found in the right ACC (*N* = 1, *p* = .008), the right insula (*N* = 1, *p* = .03), and the right STG (*N* = 1, *p* = .04), whereas visual confidence modulated the activity of the right vmPFC (*N* = 1, *p* = .001).

Beyond these, effects of stimulus modality on confidence coding were weak: for correct answers only, the amodal-confidence effect varied between auditory and visual trials in the left vmPFC (*N* = 1, *p* = .01), and between bimodal and unimodal trials in the right insula (*N* = 1, *p* = .03). The effect of auditory confidence differed as a function of the auditory presence in the right insula (*N* = 2, *p* = .002) and right OFC (*N* = 1, *p* = .03), between auditory and audiovisual trials in the left IPC (*N* = 1, *p* = .02) and the right PCC (*N* = 1, *p* = .005). Moreover, for correct auditory responses, channels in the right insula (*N* = 1, *p* = .04) and the right OFC (*N* = 1, *p* = .03) showed an interaction effect between the auditory confidence and the presence of an auditory stimulus. Finally, the effect of visual confidence differed between visual present and absent trials in the left OFC (*N* = 1, *p* = .04).

Time-locked to the response, the right insula responded to auditory confidence (*N* = 1, *p* = .02). This effect varied with stimulus modality across scattered single channels (**auditory vs audiovisual trials**: left IPC [*N* = 1, *p* = .02], left preMC [*N* = 1, *p* = .004], **presence vs absence of an auditory stimulus**: right STG [*N* = 1, *p* = .03]). While no main effect of the visual confidence was observed, its effect varied with stimulus modality in the right insula (**visual vs audiovisual trials**: *N* = 1, *p* = .02). Finally, in the left VC (*N* = 2, *p* = .04) and right dmPFC (*N* = 1, *p* = .03), the amodal confidence effect was modulated by the participant’s judgment (i.e., absence or presence judgments) for correct responses.

#### 3.4.2. Supramodal Areas for Confidence

By-channel analyses showed that, time-locked to the stimulus onset, high-gamma activity in the left premotor cortex (preMC; *N* = 1, *p* = .04) covaried with amodal confidence. At the ROI level, the insula was the only region whose high-gamma activity following stimulus onset correlated with confidence, regardless of whether the stimulus was audiovisual, auditory, or visual. Time-locked to the response, a wider network was involved, including the right IFC (*N =* 3, *p* < .001), the right mid cingulate cortex (MCC; *N* = 1, *p* = .02), the right dmPFC (*N* = 3, *p* < .001), and the right preMC (*N* = 3, *p* < .001), and, for correct answers only, the left IFC (*N* = 1, *p* = .04), with latencies spanning from −490ms to 440ms around the response (Fig.4A). At the ROI level, this was reflected in the right VC from 260ms to 330ms after the response.

**Figure 4:**
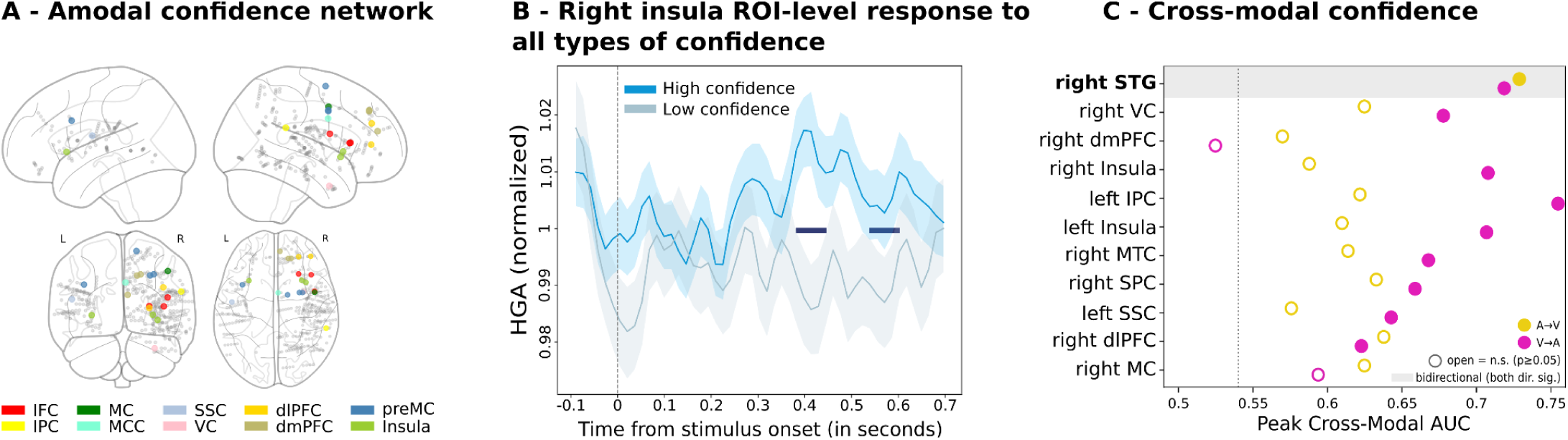
Supramodal network supporting confidence judgments. A) Supramodal confidence areas: template brain showing the regions with a significant amodal-confidence effect. B) Region-level high-gamma responses in the right insula for all types of confidence (both amodal and modality-specific); mean ± 95% CI. Note that models used confidence as a continuous variable; confidence was binarized into two equal-width bins labeled as high and low confidence only for plotting purposes. C) Cross-modal confidence decoding. Peak generalization AUC is shown for each region and transfer direction (orange, auditory→visual; pink, visual→auditory); filled symbols denote significant decoding (permutation *p* < .05), open symbols non-significant; the dashed line marks chance (AUC = 0.5), and the shaded band groups the bidirectionally-decoding region (the auditory cortex).

By-channels analysis revealed that the regions responding consistently to confidence (i.e., whether defined at the amodal level or separately for the auditory and visual responses) were the left and right insula (*N_left_* = 1, *p_left_* = .02; *N_right_*= 2, *p_right_* = .002), left and right SSC (*N_left_*= 1, *p_left_* = .009; *N_right_* = 1, *p_right_*= .02), and right MC (*N* = 1, *p* = .02). At the time around the response, it implied a wider network with an implication of the left IPC (*N* = 2, *p* = .003), right ACC (*N* = 2, *p* < .001), right IFC (*N* = 3, *p* < .001), right Insula (*N* = 1, *p* = .03), right MC (*N* = 1, *p* = .03), right MCC (*N* = 1, *p* = .002), right OFC (*N* = 1, *p* = .03), right STG (*N* = 2, *p* < .001), right dlPFC (*N* = 2, *p* = .002), right dmPFC (*N* = 4, *p* < .001), and right preMC (*N* = 2, *p* <.001), with latencies spanning from −490ms to 440ms around the response. At the ROI level, the insula was still the only region whose high-gamma activity following stimulus onset consistently responded to confidence (i.e., whether amodal or modality-specific) (Fig.4B). However, still at the ROI level, both the auditory and visual confidence recruited the auditory cortex.

Training a decoder across all trial types (i.e., auditory, visual, and audiovisual) for modality-specific confidence in one modality and testing it on the other (e.g., trained on the visual confidence and tested on the auditory confidence), we found that auditory confidence could be decoded in the bilateral insula (**left**: *AUC peak:* 0.71 at 430ms; **right**: *AUC peak:* 0.71 at 430ms), as well as in the left IPC (*AUC peak*: 0.69 at 30ms), left SSC (*AUC peak*: 0.64 at 170ms), right dlPFC (*AUC peak*: 0.62 at 470ms), right MTG (*AUC peak*: 0.67 at 330ms), right SPC (*AUC peak*: 0.66 at 610ms), right STG (*AUC peak*: 0.72 at 350ms), and the right VC (AUC peak: 0.68 at 70ms). Conversely, trained on auditory confidence, visual confidence was decoded in the right STG (*AUC peak*: 0.73 at 430ms), suggesting that the STG and visual cortex encoded both the auditory and visual confidence. This shows that sensory regions, despite exhibiting a specialization for their preferred modality at the perceptual level, also play a role in confidence, regardless of modality (Fig.4C).

## 4. Discussion

In this preregistered study, we tested whether the neural markers of detection and confidence are modality-specific or supramodal by recording high-gamma activity intracortically in patients with focal epilepsy while they performed an audiovisual near-threshold detection task. Combining intracranial recordings with single-channel, ROI-level, and multivariate analyses, we found that detection was supported by a broadly distributed supramodal network, including the bilateral insula, inferior frontal, somatosensory, and auditory cortices, with only sparse evidence for modality-specific coding. Confidence relied on a more mixed architecture that combined modality-specific and supramodal responses. Interestingly, sensory regions encoded confidence information beyond their preferred modality: high-gamma activity in the auditory cortex decoded visual confidence, with a comparable effect in the visual cortex. Together, these results suggest that detection and, to some extent, confidence involve shared cortical markers across senses, and that cortices traditionally considered to be sensory-specific participate in supramodal processing.

### 4.1. A Distributed Supramodal Detection Network

Stimulus detection was supported by a widespread supramodal cortical network. Univariate analyses identified accuracy-related high-gamma responses independent of stimulus modality, and multivariate decoding confirmed that a classifier trained to distinguish hits from misses in one modality could decode detection accuracy in the other. This cross-modal generalization corroborates and extends previous scalp-level findings of supramodal detection signatures (Sanchez et al., 2020; Niedernhuber et al., 2022; Noel et al., 2018). By localizing bidirectional cross-modal generalization to the dorsomedial prefrontal cortex, insula, inferior parietal cortex, and superior temporal gyrus, our results point to a shared supramodal detection network involving several cortical areas. Our predominantly supramodal effects contrast with modality-specific detection signatures previously reported at the scalp level (Koivisto & Revonsuo, 2010; Eklund & Wiens, 2019; Filimonov et al., 2022; Ciupińska et al., 2025); however, we note that our recordings may have missed them, particularly in the visual modality, as they predominantly sampled associative rather than early visual regions (see Limitations). Coverage is a less plausible explanation for audition: the STG was well sampled but yielded only one channel distinguishing auditory hits from misses, suggesting a genuine but comparatively weak modality-specific signal.

We also examined whether cortical activity differentiated participants’ judgments of presence and absence. Differences, both irrespective of accuracy and for correct responses only, were concentrated in frontal regions, notably the orbitofrontal, the inferior frontal, and dorsolateral prefrontal cortices. These results were consistent with previous work dissociating the correlates of detection from those of presence vs absence judgment (Mazor et al., 2020). This dissociation has been interpreted as corroborating the involvement of additional, higher-order, processes for absence judgments, mostly localized in the frontal cortex (Mazor & Fleming, 2020; Mazor et al., 2020; Mazor et al., 2022).

### 4.2. A Supramodal Code for Confidence

Although confidence was associated with modality-specific responses in regions that differentiated auditory from visual confidence (the right motor cortex and right superior temporal gyrus), a supramodal structure emerged: at the ROI level, the insula was the only region whose high-gamma activity following stimulus onset tracked confidence, irrespective of whether it was reported on the amodal, auditory, or visual scale. A broader supramodal network appeared when activity was locked to the response, including the insula, inferior frontal cortex, dorsomedial prefrontal cortex, anterior and mid-cingulate cortices, and inferior parietal cortex. Applying cross-modal decoding to confidence, a decoder trained on visual confidence responses could predict auditory confidence responses in several regions, including the bilateral insula, superior temporal gyrus, inferior parietal cortex, and visual cortex. The recurrent involvement of the inferior frontal cortex in our supramodal confidence effects converges with fMRI evidence that the right inferior frontal gyrus integrates confidence across memory and perception (Morales et al., 2018), as well as independently of sensory modality during probabilistic inference (Meyniel & Dehaene, 2017), suggesting that its amodal role in confidence generalizes across tasks as different as learning and detection.

This provides, to our knowledge, the first evidence for cross-modal generalization of a neural code for confidence during detection. Previous studies reported that metacognitive performance correlates across modalities, and that confidence-related electrophysiological responses show no interaction with modality (Faivre et al., 2018). Our findings extend this to a detection task, showing that confidence can be actively decoded across modalities, consistent with a shared representation of confidence across the senses. However, two considerations temper this interpretation. First, the modest number of participants and channels contributing to each analysis warrants caution. Second, because confidence covaries with decision time, a decoder trained on confidence could capture a modality-independent decisional signal, such as a shared reaction-time correlate, rather than a confidence representation per se. We addressed this directly by regressing the detection reaction time from the high-gamma activity at every channel and time point, fitting the nuisance regression within each modality, and then repeating the cross-modal decoding on the residualised activity. We found that cross-modal transfer was essentially unchanged (see SI.3.4). Moreover, we asked whether reaction time was decodable in the same regions and time windows: a decoder trained to separate fast from slow trials generalized across modalities only modestly, below the confidence and detection transfer it was meant to explain (see SI.3.4). Together, these analyses indicate that the cross-modal generalization of both detection and confidence is not an artifact of a shared reaction-time signal.

### 4.3. Specialized Sensory Regions Encode More than Perceptual Information

We replicated the implication of specialized sensory cortices for multisensory perception, both in the auditory and visual cortex (Fiorini et al., 2021; Noesselt et al., 2010; Ferraro et al., 2020). In addition to this multisensory perceptual processing, we observed that the auditory cortex encoded detection processing irrespective of the stimulus modality. Moreover, sensory cortices carried neural activity that encoded confidence beyond their preferred modality: high-gamma activity in the auditory cortex (superior temporal gyrus) decoded both visual and auditory confidence, with a parallel effect observed in the visual cortex for auditory confidence. It echoes reports that confidence is decodable from widely distributed cortical networks, including regions not classically associated with metacognition (Nakuci et al., 2024; Goueytes et al., 2025; Gherman et al., 2024), and suggests that the involvement of sensory cortices in confidence is not confined to its preferred input.

### 4.4. The Insula as a Candidate Integrative Region

Finally, across analyses, the insula recurred as one of the most consistent contributors to both detection and confidence irrespective of the sensory modality, echoing its role in evidence accumulation, decision-making, and confidence evidenced in prior work (Pereira et al., 2020; Goueytes et al., 2025; Lauriers et al., 2026; Yeon et al., 2020). These effects were not spatially homogeneous: detection- and confidence-related responses tended to arise at more anterior insular sites, consistent with a decisional or monitoring role, whereas some auditory-related responses arose more posteriorly, possibly reflecting proximity to the auditory cortex (see SI.3.5).

### 4.5. Limitations

There are several limitations to our conclusions. First, as in all intracranial studies, electrode placement was determined solely by clinical considerations, resulting in uneven and incomplete coverage; the early visual cortex, in particular, was sparsely sampled due to implantation constraints, which likely contributed to the scarcity of modality-specific visual effects. Moreover, our sample was small, largely due to clinical constraints and limited access to patients capable of performing our demanding psychophysical task. As a result, several ROI-level and decoding analyses relied on regions contributed by only one or a few participants, limiting the robustness and generalizability of the corresponding effects. Our decoding results show that a common signal predicts detection and confidence across modalities, but cannot establish whether this signal is computed locally within each sensory cortex or instead reflects top-down feedback from a supramodal hub such as the insula. Both accounts describe a supramodal architecture, but only the former implies that sensory cortices themselves perform amodal computations. Finally, because amodal and modality–specific confidence were rated sequentially on every trial and were coupled behaviorally, we cannot fully exclude the possibility that participants generated a partly unitary confidence signal redistributed across the three measures (see Figure S1 for correlation between the visual and auditory confidences). Consequently, cross-modal confidence decoding may partly reflect this shared upstream judgment rather than fully distinct modality-specific confidence codes converging in the supramodal cortex.

### 4.6. Conclusions

Our results revealed that detection involves a supramodal network, distributed across cortical regions. Moreover, they indicate that even specialized sensory cortices participate in supramodal processing, not only for detection, but also at the confidence level. Together, these findings favor a neural architecture drawn on largely shared modality-general cortical resources broadly distributed across the brain at both detection and confidence levels.

## Contributions

PP: conceptualisation, methodology, software, formal analysis, investigation, data curation, writing – original draft, writing – review & editing, visualization; AR: resources, writing – review & editing; FB: Software, investigation, data curation, writing – review & editing; CD: conceptualization, software, writing – review & editing; DG: software, data curation, writing – review & editing; SM: formal analysis, writing – review & editing; MP: methodology, writing – review & editing; DH: resources, writing – review & editing; PK: resources, writing – review & editing; LM: resources, writing – review & editing; JCG: methodology, investigation, data curation, writing – review & editing; LG: conceptualization, methodology, validation, visualisation, supervision, writing – review & editing; NF: conceptualisation, methodology, software, validation, formal analysis, resources, data curation, visualisation, supervision, funding acquisition, writing – original draft, writing – review & editing

## Acknowledgments

Funded by the European Union. Views and opinions expressed are however those of the author(s) only and do not necessarily reflect those of the European Union or the European Research Council Executive Agency. Neither the European Union nor the granting authority can be held responsible for them. This work is supported by an ERC grant (Volta, 101125379) awarded to NF.

## Declaration of generative AI and AI-assisted technologies in the manuscript preparation process

During the preparation of this work, AI-assisted technologies were used to improve language (as English is not the native language of the authors) and to double-check for coding errors.

