## supplementary information for "Supramodal Coding of Detection and Confidence in the Human Cortex"

**Table S1**: Demographic Information of Included Patients.
To preserve patients' anonymity, ages are reported in a class format

| Patient | Gender | Age | Dominant Hand | Number of blocs performed |
|---|---|---|---|---|
| 1 | M | 20-29 | Both | 6 |
| 3 | F | 20-29 | Right | 6 |
| 4 | M | 20-29 | Right | 6 |
| 6 | M | 30-39 | Right | 4 |
| 8 | M | 20-29 | Right | 3 |
| 10 | M | 40-49 | Right | 4 |
| 11 | M | 20-29 | Right | 6 |
| 12 | M | 40-49 | Right | 6 |

**Table S2**: Participants included in each neural model

| | Number of participants excluded | Exclusion reason |
|---|---|---|
| **Analysis locked on the stimulus onset** | | |
| ***Amodal analysis*** | | |
| ~ modality | 0 | / |
| ~ accuracy | 1 | Lack of incorrect responses |
| ~ modality * accuracy | 1 | Lack of incorrect responses |
| ~ confidence | 2 | Lack of variation in confidence judgment |
| ~ confidence * modality | 2 | Lack of variation in confidence judgment |
| ***Auditory analysis*** | | |
| ~ accuracy * modality | 3 | No incorrect responses when the auditory stimulus is absent |
| ~ confidence * modality | 2 | Lack of variation in confidence judgment |
| ~ confidence * modality (correct resp. only) | 3 | Lack of variation in confidence judgment |
| ***Visual analysis*** | | |
| ~ accuracy * modality | 4 | No incorrect responses when the visual stimulus is absent |
| ~ confidence * modality | 2 | Lack of variation in confidence judgment |
| ***Link between scales*** | | |
| ~ accuracy | 0 | / |
| ~ confidence | 2 | Lack of variation in confidence judgment |
| ***Multivariate analysis*** | | |
| Detection | 0 | / |
| Confidence | 1 | binning the confidence ratings resulted in only one confidence level for correct responses, leading to the exclusion of the following regions: left MTG and left STG. |
| **Analysis locked on the response** | | |
| ***Amodal analysis*** | | |

| | | |
|---|---|---|
| ~ modality | 0 | / |
| ~ accuracy | 0 | / |
| ~ judgment | 0 | / |
| ~ modality * accuracy | 4 | No incorrect presence judgments |
| ~ confidence | 2 | Lack of variation in confidence judgment |
| ~ confidence * modality | 2 | Lack of variation in confidence judgment |
| ~ confidence * judgment | 2 | Lack of variation in confidence judgment |
| ~ confidence * accuracy in absence judgments | 2 | Lack of variation in confidence judgment |
| ***Auditory analysis*** | | |
| ~ accuracy * modality | 3 | No incorrect responses when the auditory stimulus absent |
| ~ confidence * modality | 2 | Lack of variation in confidence judgment |
| ***Visual analysis*** | | |
| ~ accuracy * modality | 2 | No incorrect responses when the visual stimulus is absent |
| ~ confidence * modality | 0 | / |
| ***Link between scales*** | | |
| ~ accuracy | 0 | / |
| ~ confidence | 2 | Lack of variation in confidence judgment |

#### SI 1 Individual ROIs for Multivariate Analyses

- right dmPFC (of one participant)
- left IPC (of two participants)
- left Insula (of one participant)
- right Insula (of one participant)
- left SSC (of one participant)
- right dlPFC (of one participant)
- right MC (of one participant)
- right STG (of three participants)
- left STG (of one participant)
- right VC (of two participants)
- left MTG (of one participant)
- right MTG (of one participant)
- right SPC (of one participant)

### SI 2. Regrouping of MarsAtlas ROIs

- **Inferior Parietal Cortex (IPC):** dorsal inferior parietal cortex, ventral inferior parietal cortex
- **Medial Parietal Cortex (MPC):** medial parietal cortex
- **Superior Parietal Cortex (SPC)**: superior parietal cortex, medial superior parietal cortex
- **Somatosensory Cortex (SSC)**: ventral somatosensory cortex, dorsolateral somatosensory cortex, dorsomedial somatosensory cortex
- **Premotor Cortex (preMC)**: dorsolateral premotor cortex, dorsomedial premotor cortex
- **Motor Cortex (MC)**: ventral motor cortex, dorsolateral motor cortex, dorsomedial motor cortex
- **Insula**: Insula
- **Inferior Frontal Cortex (IFC):** rostral ventrolateral prefrontal cortex, rostral ventral premotor cortex
- **Orbitofrontal Cortex (OFC):** ventral orbitofrontal cortex, ventrolateral orbitofrontal cortex, ventromedial orbitofrontal cortex
- **dorsolateral Prefrontal Cortex (dlPFC):** rostral dorsolateral inferior prefrontal cortex, rostral dorsolateral superior prefrontal cortex, caudal dorsolateral prefrontal cortex
- **dorsomedial Prefrontal Cortex (dmPFC)**: caudal dorsomedial prefrontal cortex, rostral dorsal prefrontal cortex
- **ventromedial Prefrontal Cortex (vmPFC):** ventromedial prefrontal cortex, prefrontal medial cortex
- **Superior Temporal Gyrus (STG; i.e., auditory cortex):** rostral superior temporal cortex, caudal superior temporal cortex
- **Middle Temporal Gyrus (MTG):** rostral middle temporal cortex, caudal middle temporal cortex
- **Visual Cortex (VC):** rostral inferior temporal cortex, medial inferior temporal cortex, lateral visual cortex, rostral medial visual cortex, cuneus, superior visual cortex, caudal medial visual cortex
- **Isthmus Cingulate Cortex (ICC)**: isthmus cingulate cortex
- **Anterior Cingulate Cortex (ACC):** anterior cingulate cortex
- **Mid Cingulate Cortex (MCC):** mid cingulate cortex
- **Posterior Cingulate Cortex (PCC)**: posterior cingulate cortex

**Table S3:** Median reaction times in seconds (*with interquartile range*) as a function of the condition of presentation and the response.

| **Experimental condition** | **Judge Absent** | **Judge Present** |
|---|---|---|
| AV | 2.3 *(1.09)* | 1.87 *(0.92)* |
| A | 2.09 *(0.83)* | 1.97 *(0.92)* |
| V | 2.26 *(1.11)* | 1.87 *(0.94)* |
| Absent | 2.09 *(0.9)* | 2.22 *(1.01)* |

### SI 3: Additional Neural Results

#### SI 3.1. Neural Correlates of Perception

By-channel analysis showed that other areas than the auditory and visual cortex had a differentiated activity between auditory and visual trials including the left MTG ($N$ = 2, $p$ <.001), the left SSC ($N$ = 1, $p$ = .04), the right MC ($N$ = 3, $p$ = .002), the right SPC ($N$ = 2, $p$ = .003), other channels survived FDR correction but with a prevalence not exceeding chance by permutation ($N$ = 1 in the right IFC and $N$ = 1 in the right IPC).

Single-channels multimodal evoked responses (i.e., difference in stimulus-evoked responses between bimodal and unimodal trials) showed that channels that survived FDR correction but with a prevalence not exceeding chance by permutation were localized in the right VC ($N$ = 1), right IPC ($N$ = 1), and the right IFC ($N$ = 1).

When focusing only on correct responses, differences between auditory and visual trials were observed in the left and right IFC ($N_{left}$ = 1, $p_{left}$ = .005; $N_{right}$ = 1, $p_{right}$ = .02), the left MTG ($N$ = 1, $p$ = .01), the left SSC ($N$ = 1, $p$ = .006), the right MC ($N$ = 2, $p$ < .001), the right STG ($N$ = 2, $p$ = .002), the right VC ($N$ = 2, $p$ < .001); and between bimodal and unimodal trials in the right VC ($N$ = 1, $p$ = .006).

We also investigated the remaining neural correlates of perception at the time around the responses. By-channel analysis showed a differentiated activity between auditory and visual trials in the left MTG ($N$ = 3, $p$ < .001) the right IFC ($N$ = 1, $p$ = .04), with one channel in the right IPC, two channels in the right VC and one channel in the right STG not exceeding chance-level by permutation; and between bimodal and unimodal trials in the right IPC ($N$ = 1, $p$ = .04) and the right VC ($N$ = 4, $p$ = .004). Focusing on correct answers, differences between auditory and visual trials were observed in the left SSC ($N$ = 2, $p$ = .01) and the right IPC ($N$ = 1, $p$ = .02), and between bimodal and unimodal trials in the right dmPFC ($N$ = 1, $p$ = .009).

##### Presence vs Absence of a Stimulus

At the time around the response, we additionally performed by-channel analysis to investigate areas where high-gamma responses differentiated absent and present trials. Such effects were observed in the left OFC ($N$ = 2, $p$ = .004), the left and right VC ($N_{left}$ = 1, $p_{left}$ = .03; $N_{right}$ = 9, $p_{right}$ <.001), the right IFC ($N$ = 1, $p$ = .03), the right SPC ($N$ = 5, $p$ < .001) and the right preMC ($N$ = 1, $p$ = .02), with latencies spanning from -480ms to 280ms around the response, channels in the right IPC ($N$ = 1) and the right insula ($N$ = 1) did not exceed chance by permutation. These effects were also observed in the left VC ($N$ = 1, $p$ = .009) and right dlPFC ($N$= 1, $p$ = .02) when focusing on correct responses.

Moreover, evoked high-gamma responses differentiating presence and absence judgments that survived FDR correction but with a prevalence not exceeding chance by permutation were located in the left VC ($N$ = 1), the left dlPFC ($N$ = 1), the right insula ($N$ = 1), the right MPC ($N$ = 1), the right MTG ($N$ = 1), and the right vmPFC ($N$ = 1).

#### SI 3.2. Neural Correlates of Detection

Focusing specifically on amodal responses, channels that survived FDR correction but with a prevalence not exceeding chance by permutation were localized in the left SSC ($N$ = 1), the left dlPFC ($N$ = 1), the right IFC ($N$ = 1), the right PCC ($N$ = 2), the right SPC ($N$ = 1), the right SSC ($N$ = 1), the right VC ($N$ = 1) and the right vmPFC ($N$ = 1).

Turning to modality-specific effects, at around stimulus onset, channels in the right OFC ($N$ = 1, $p$ = .02) and right SPC ($N$ = 1, $p$ = .02) showed differentiated activity as a function of visual accuracy between visual-present and visual-absent trials.

##### Response-locked Results: Detection Effects

Analysis at the time around the response showed that multiple regions reflected stimulus detection irrespectively of its nature (i.e., amodal accuracy and modality-specific auditory and visual accuracy), including the left and right insula ($N_{left}$ = 1, $p_{left}$ = .02; $N_{right}$ = 2, $p_{right}$ < .001), left OFC ($N$ = 1, $p$ = .01), left VC ($N$ = 1, $p$ = .02), right ACC ($N$ = 1, $p$ = .01), right IFC ($N$ = 1, $p$ = .02), right MC ($N$ = 2, $p$ < .001), right MTG ($N$ = 1, $p$ = .04), right dlPFC ($N$ = 1, $p$ = .04), and right dmPFC ($N$ = 1, $p$ = .01). Moreover, the left MTG ($N$ = 1, $p$ = .03), and the left SSC ($N$ = 1, $p$ = .01) had a differentiated activity between auditory and visual accuracy.

Focusing on amodal responses, only the right insula had a differentiated activity between correct and incorrect responses ($N$ = 2, $p$ = .02); other channels survived FDR correction but with a prevalence not exceeding chance by permutation (***left IPC:*** $N$=1, ***left insula***: $N$=1, ***left OFC***: $N$=1, ***right MC***: $N$=1, ***right OFC***: $N$=1, ***right VC***: $N$=1, ***right dmPFC***: $N$=1). In addition, three channels in the left MTG ($p$ < .001) and one channel in the right IFC ($p$ = .04) were specific to the modality, with an interaction between the accuracy of the response and the modality (visual vs auditory trials). Moreover, one channel in the right IPC ($p$ = .04) and four channels in the right VC ($p$ = .004) were considered multimodal, with an interaction between response accuracy and modality (bimodal vs unimodal trials).

For modality-specific responses, single-channels evoked high gamma responses differed between auditory correct and incorrect responses in the right insula ($N$ = 2, $p$ < .001) and right OFC ($N$ = 1; $p$ = .02). Moreover, the left and right SSC ($N_{left}$ = 1, $p$ = .008; $N_{right}$ = 3, $p$ = .004), the right IFC ($N$ = 1, $p$ = .02), the right insula ($N$ = 2, $p$ < .001), the right OFC ($N$ = 2, $p$ <.001), and the right vmPFC ($N$ = 1, $p$ = .008) showed an interaction effect between the presence or absence of an auditory stimulus and the auditory accuracy. For visual responses, the right OFC ($N$= 1, $p$ = .02) showed an interaction effect between the presence or absence of a visual stimulus and the visual accuracy.

#### SI 3.3 Neural Correlates of Confidence

For incorrect answers only (i.e., miss trials; analysis-locked on the stimulus onset), differences in HGA as a function of the amodal confidence were observed in the right ACC ($N$ = 1, $p$ = .01) and the right insula ($N$ = 1, $p$ = .04), with one channel in the right OFC not exceeding chance level by permutation.

Channels showing a difference as a function of the amodal confidence that survived FDR correction but with a prevalence not exceeding chance by permutation were localized in the left insula ($N$ = 1) the left SSC ($N$ = 1), the right insula ($N$ = 1), the right MC ($N$ = 1), and the right dlPFC ($N$ = 1). For the time around the response, they were localized in the left insula ($N$ = 1), the right IPC ($N$ = 1), the right insula ($N$ = 1), the right VC ($N$ = 1), and the right dlPFC ($N$ = 1).

#### SI 3.4 Exploratory Decoding Results with Reaction Times

We performed additional, exploratory, decoding analyses to test whether our cross-modal results, both at the detection and confidence levels, could be an artefact from the shared reaction-time signal. These analyses reused the cross-modal decoding pipeline described in the main text.

##### Regressing out Reaction Time.

For each channel and time point, high-gamma activity was regressed on reaction time within each modality using ordinary least squares, and the residuals replaced the original activity before decoding. Because the decoded label (hit versus miss, or high versus low confidence) was never included in this regression, only the component of high-gamma activity that was collinear with reaction time was removed; label-related variance orthogonal to reaction time was preserved. Fitting the regression within modality prevented removal of cross-modal structure and was applied identically to training and test trials.

Removing the reaction-time-collinear component of high-gamma activity left cross-modal decoding largely intact. Among regions with significant detection transfer in the main analysis ($n$ = 14), the peak AUC fell only from a mean of 0.75 to 0.71 after residualization. Among regions with significant confidence transfer ($n$ = 10), peak AUC was essentially unchanged (mean 0.69 before, 0.69 after). Assessed against the same 1000-permutation null as the main decoding, 13 of 14 detection effects and 8 of 10 confidence effects remained significant ($p < .05$) after residualization.

##### Decoding Reaction Time

To test whether reaction time was itself decodable where confidence and detection generalised, trials were labelled fast or slow by a within-modality median split of reaction time, and the detection decoding was repeated with this label (training in one modality, testing in the other). For the confidence design, in which training and test are two reports of the same trials (as the auditory and visual confidence are given on each trial), reaction-time decodability was assessed within-sample by cross-validation against a label-permutation null.

Reaction times were only modestly decodable across modalities: for the detection-design, only 5 regions were found to be able to decode RTs (against 14 for accuracy), including the left insula, the right SPC, the right VC, the left SSC, and the right STG. The same was observed for the confidence-design, with, again only 6 regions with significant decoding results, including the left insula, the left SSC, the right dlPFC, the right STG, the right MTG, the right SPC (against 10 in the main cross-modal confidence pipeline).

Together, the residualization and reaction-time-decoding analyses indicate that cross-modal generalization of detection and confidence is not an artifact of a shared decisional or reaction-time signal.

#### SI 3.5 Insular Results: Exploration of Anterior vs Posterior Insula

Given our limited sample size and the fact that the MarsAtlas parcellation does not distinguish the anterior and posterior insula, our main analyses considered the insula as a single region. Nevertheless, for each of our significant by-channel effects, we explored whether the channels were localized to the anterior or posterior part of the insula. We determined the localization of channels to the anterior or posterior insula through direct visualization of each patient's individual anatomical scans. Channels showing amodal, supramodal effects – including amodal stimulus detection, accuracy irrespective of modality, and confidence irrespective of type – were more frequently localized to the anterior insula (3/4 channels for amodal detection, 2/3 for accuracy irrespective of type, 2/3 for confidence irrespective of type). Conversely, channels showing auditory-specific confidence effects were predominantly localized to the posterior insula (2/2 for auditory confidence; 1/1 for auditory confidence restricted to correct trials; 2/2 for auditory confidence between present and absent stimuli). This pattern was consistent across both stimulus-locked and response-locked analyses, raising the possibility of a functional gradient within the insula, whereby more anterior sites contribute to supramodal, amodal representations of detection and confidence, while more posterior sites are preferentially engaged in auditory-specific processing. However, given the very small number of channels involved in each contrast, this dissociation is only exploratory.

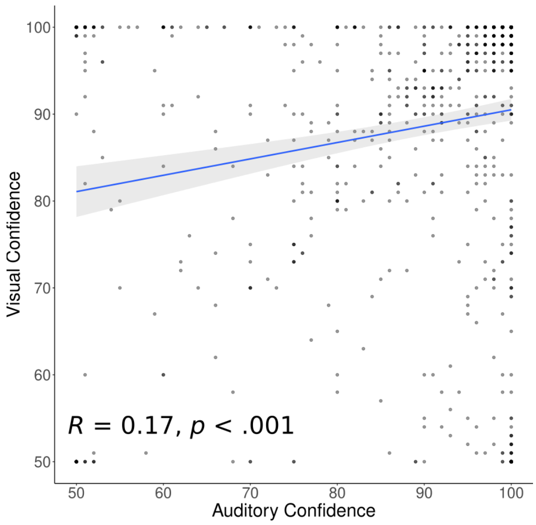


**Figure S1**: Correlation between the visual and the auditory confidence, when the amodal detection response is correct and the stimulus present.
